# Theranostic cranial implant for hyperspectral light delivery and microcirculation imaging without scalp removal

**DOI:** 10.1101/720599

**Authors:** Nami Davoodzadeh, Mildred S. Cano-Velázquez, Carrie R. Jonak, David L. Halaney, Devin K. Binder, Juan A. Hernández-Cordero, Guillermo Aguilar

## Abstract

Light based techniques for imaging, diagnosing and treating the brain have become widespread clinical tools, but application of these techniques is limited by optical attenuation in the scalp and skull. This optical attenuation reduces the achievable spatial resolution, precluding the visualization of small features such as brain microvessels. The goal of this study was to assess a strategy for providing ongoing optical access to the brain without the need for repeated craniectomy or retraction of the scalp. This strategy involves the use of a transparent cranial implant and skin optical clearing agents, and was tested in mice to assess improvements in optical access which could be achieved for laser speckle imaging of cerebral microvasculature. Combined transmittance of the optically cleared scalp overlying the transparent cranial implant was as high as 89% in the NIR range, 50% in red range, 24% in green range, and 20% in blue range. *In vivo* laser speckle imaging experiments of mouse cerebral blood vessels showed that the proposed optical access increased signal-to-noise ratio and image resolution, allowing for visualization of microvessels through the transparent implant, which was not possible through the uncleared scalp and intact skull.

## INTRODUCTION

From the deep blue to the near-infrared (NIR) is the region among the wide spectrum of electromagnetic waves that we know as light. This delicate region has a distinct advantage for interrogating biological tissue due to its unique photon energy range of 0.5–3 eV. At energies above this range, C–C and C–H bonds dissociate and ionization can occur, while at energies below this range, water absorption dominates the transmittance, preventing any specific targeting of molecules^1^. The noninvasive interactions with tissue of photons within this energy range have been used in numerous approaches to assess brain health and treat its diseases. Some of these treatment practices have been used in clinical applications such as photobiomodulation (for wound healing, tissue repair, anti-inflammatory therapy^2^, and chronic traumatic brain injury^3^) and photodynamic therapy (PDT) which is now clinically approved to treat brain cancer^4^. Optical dissection of brain circuits methods, known as optogenetic therapies, have resulted in the treatment of Parkinson^5^, cocaine addiction^6^, depression^7^, chronic pain^8^, and laryngeal paralysis^9^ in animals and may one day be translated for use in humans. Consequently, bio-optical technologies for adequate delivery of light to the brain have been rapidly advancing^10^. The poor penetration depth of light, particularly in the visible range, has limited the clinical applications of photosensitizer activation-based methods. For instance, PDT is currently only used for the treatment of superficial lesions, or of lesions accessible during surgery or through endoscopy^4^.

Light-based techniques have also been used for brain health assessment. Numerous methods have been widely developed and applied in various brain disease diagnostics. Imaging of brain microcirculation (i.e. blood flow in vessels <150 μm in diameter), which plays a critical role in the physiological processes of tissue oxygenation and nutrient exchange, has enabled us to noninvasively extract morphological and functional information^11,12^. Morphological information provides vessel density, perfusion rate, vessel diameter, and dynamic measurements of microcirculatory blood flow velocity and blood cell concentration^11^, while functional data provides information on blood oxygenation, changes in metabolism, regional chemical composition, etc^12^. Various imaging modalities have been developed which are capable of measuring blood flow. Although they have resulted in a number of discoveries, there are notable drawbacks which should be considered in developing a scalable and real-time method for routine and long-term brain microcirculation monitoring. Indocyanine green video angiography (ICGVA) requires injection of dyes and cannot provide continuous assessment of vessel perfusion^13,14^. Optical coherence tomography (OCT)^15^, photoacoustic imaging (PAM)^16^, and fluorescence microscopy approaches like confocal microscopy or the two-photon variant^17^ are based on laser scanning, which is limiting to the temporal resolution and field of view^18^, although recent technical improvements in OCT and PAM have been reported which provide sufficient temporal resolution for epileptic seizure monitoring^19–21^. Fast full-field laser Doppler perfusion imaging (ffLDPI)^17^ cannot simultaneously provide microvascular structural and functional information due to lack of adequate spatiotemporal resolution^22^.

Analyzing intensity fluctuations in scattered laser light, i.e. speckles, has been reported to provide useful biological information. Laser Doppler, photon correlation spectroscopy, and time-varying speckle operate by analyzing speckles from a single point in the flow field and thus require some form of scanning to produce velocity maps, adding time to the procedure^17^. Laser speckle imaging (LSI), on the other hand, is a full-field, real-time, noninvasive, and noncontact imaging method which is sensitive to both the speed and morphological changes of the scattering particles, and is capable of mapping relative velocity in flow fields such as capillary blood flow^17^. By utilizing spatio-temporal statistics of speckle, LSI produces blood flow maps in a fraction of a second without the need for scanning, making it a true real-time technique^17^. Like LSI, ffLDPI can provide full field information in a short time, but LSI has the advantage of being relatively simple and inexpensive since it does not require a high-speed imaging array. This make LSI a scalable and compatible modality which has been used in wearable and routing brain monitoring applications^23^, solely or coupled with other imaging modalities^17^.

Proving theranostic optical access to the brain for health assessment is complicated due to the static and dynamic effects of optical scattering and absorption by the inhomogeneous skull and scalp tissues. These effects drastically limit the achievable imaging field of depth and resolution. Therefore, brain imaging has been predominantly demonstrated in rodent open-skull and thinned skull models. Likewise, in our own prior studies we have found that optical imaging through the intact skull of mice diminishes detection of the intrinsic optical signals^24–27^. Moreover, the optical properties of the inhomogeneous cranial bone overlap with those of brain tissue and jointly these factors decrease the accuracy and reliability of the data. Studies in the literature have addressed this optical barrier with various approaches including imaging the brain directly with an opened skull^28^, thinned and polished skull preparations^29,30^, temporary optical clearing of the skull using optical clearing agents^31–33^, and implanting glass or PDMS windows^34,35^. Glass and PDMS windows are powerful research techniques, but are not appropriate for human application as permanent cranial implants for patients. Like skull thinning and polishing techniques, glass-based windows compromise protection for the brain due to the extremely low fracture toughness of typical glasses (KIC = 0.7-0.9 MPa m1/2)^36^ which increases potential for catastrophic failure by fracture, while the effect of skull optical clearing agents for long-term use on human skull is still unknown^24^. A number of biomedical considerations including biocompatibility, mechanical strength, and ageing should be examined in order to create an optical window for eventual clinical application^24^. Conventional cranial prosthesis including titanium, alumina, and acrylic^37^, have not provided the requisite combination of transparency and toughness required for clinically-viable transparent cranial implants. To address this challenge, our group has previously introduced a transparent nanocrystalline yttria-stabilized-zirconia cranial implant material, which possesses the mechanical strength and biocompatibility which are prerequisites for a clinically-viable permanent cranial implant for patients^24–27,38–46^. This implant has been referred to in the literature as the “Window to the Brain” (WttB) implant. Figure 1 shows an illustration of implant placement envisioned in eventual human application providing theranostic optical access for light delivery and high resolution and chronic brain monitoring without scalp removal.

**Figure 1.**
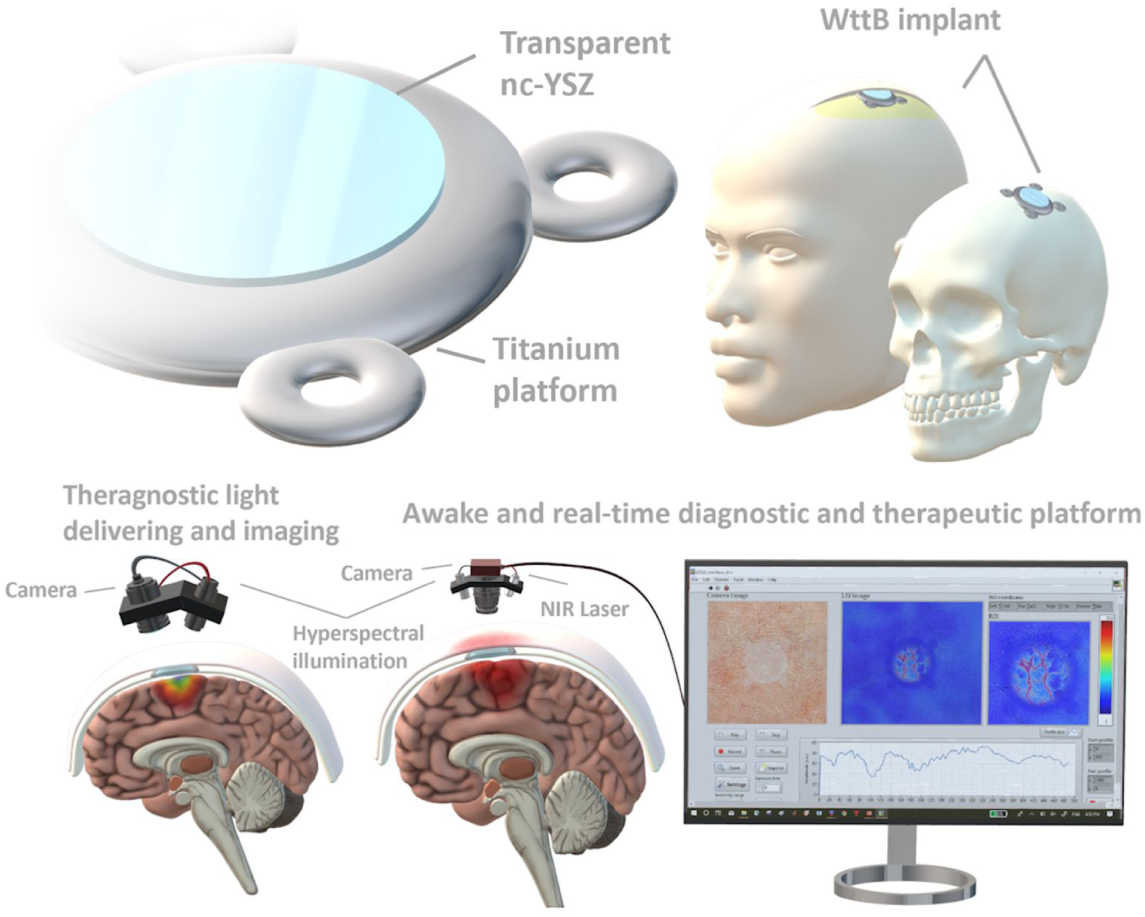
Illustration of the Window to the Brain cranial implant concept. The implant will be attached to the skull beneath the scalp, to allow for optical transmission to and from the brain.

Because this cranial implant is sought as a means to obtain optical access for post-operative and prolonged theranostic purposes without scalp removal, scalp scattering must be overcome. Therefore, in this current study we evaluate the use of optical clearing agents (OCAs) in the scalp of mice. As shown in previous reports, OCAs provide greater optical probing depths and better contrast, as well as improved light focusing and spatial resolution^47–49^. In this report, the enhancement in optical access to the brain using scalp optical clearing and WttB implant is evaluated through two experiments of comparing *ex vivo* optical transmittance measurements (Experiment 1) and *in vivo* brain hemodynamics imaging (Experiment 2) in mice.

## RESULTS

### Experiment 1: Ex vivo characterization of light delivery

Figure 2(a) illustrates the spectrotemporal effect of OCA on scalp. Absolute change in the optical transmittance spectra (referenced to baseline before applying OCA) for up to 50 minutes after applying OCA is shown in Figure 2(a). A primary change in optical transmittance is noticeable immediately upon applying the OCA (0 minutes). A sudden increase then occurs after 10 minutes which then gradually decreases. The curved black lines on the 3D figure depict transparency increases by 3.5%. A maximum optical transmittance of 90% (41% increase above baseline values) was obtained after 11 minutes from applying OCA for the wavelength of 889 nm. The upper 2D map in the figure also reflects the spectrotemporal behavior of the optical cleared scalp (OCS), showing the ideal time to optically access the brain across the OCS. As an example, the dashed line represents the temporal change at a wavelength of 810 nm (the wavelength used for LSI in Experiment 2). A time-window of ∼13 minutes for using the OCS was evaluated, in which the OCS transparency (at 810 nm) is ∼36% higher than the untreated scalp tissue (Figure 2(b)). The top 3% of optical transmittance improvement was achieved within minutes 5 to 18 after applying OCA, associated with an absolute change from 33% up to 36%. Figure 2(c) shows the temporal change in optical properties of the scalp tissue from the moment OCA was applied (0 minute) up to 40 minutes after application. A higher light transmission and transparency is noticeable at minute 10 which gradually decreases after the 20-minute mark.

**Figure 2.**
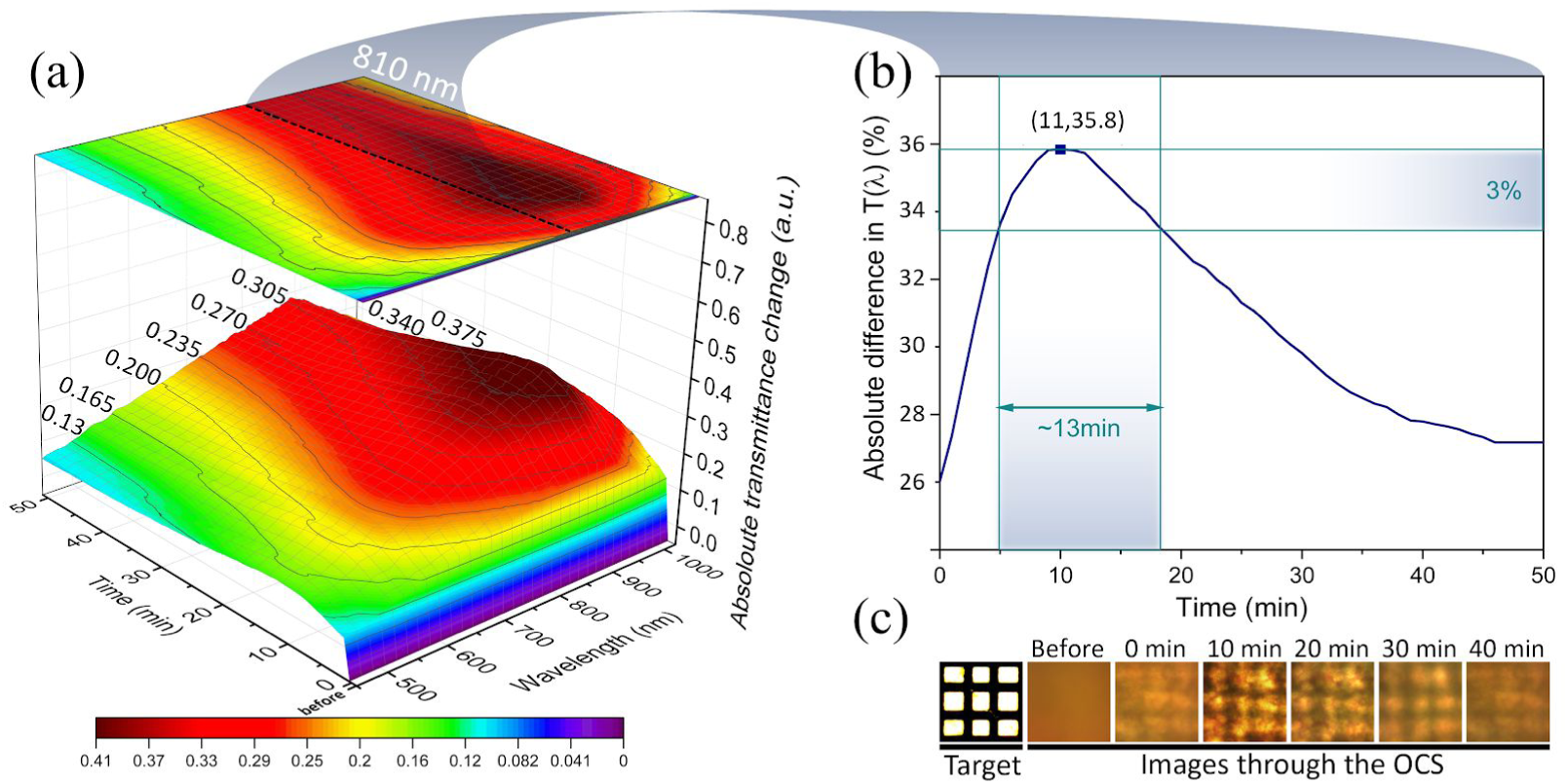
Temporal effect of OCA on scalp (Experiment 1). a) Spectrotemporal behavior of OCS, absolute change in optical transmittance is shown versus wavelength and time. b) Temporal change in optical transmittance at a wavelength of 810 nm. A time-window of 13 minutes is augmented to the plot showing the duration of time transmittance is highest in the OCS. c) Visual comparison of photographs of a resolution target captured through scalp tissue before, immediately after (0 minute), and up to 40 minutes after applying OCA. The distance between the lines in the target is ∼150 μm.

A set of photographs of an NBS 1963A resolution target imaged through the implant, skull, and scalp provides a visual comparison of the light transmittance and image quality over the samples (Figure 3(a)). Optical transmittance of skin, skull, and implant are shown in Figure 3(b). All three of the scalp, skull, and implant show an increasing trend in transmittance as the wavelength increases. The implant has higher optical transmittance compared to the skull in the range of 450-820 nm. In the range of 525 - 600 nm, skull shows a drop in the transmittance which was not observed in the implant spectra due to the high. This lower optical transmission is due to the higher optical absorption of hemoglobin in that range ^50,51^. The same change in the slope at 525 - 600 nm, appeared in the scalp tissue as well. A comparison between the mouse scalp and skull optical transmittance shows that optical extinction by the scalp is notably higher than that of the skull, revealing that the scalp is the primary limiting factor here. The optimum effect of OCA was considered for the comparisons in Figure 3(c), 3(d), 3(e), and (f)). Figure 5(c) shows an 11% improvement (gray dashed line) in optical transmittance at 450 nm in OCS which increases up to 41% (at 889 nm) at longer wavelengths. Optical transmittance of the scalp+skull stack is shown in Figure 3(d) (dark green line). As expected, it resulted in a similar optical transmittance to the scalp (limited by the high optical extinction in the scalp). In OCS+skull, the optical transmittance increased (gray dashed line) by 2.3% (at 450 nm) up to 22% (at 889 nm). The scalp+implant stack also has a similar optical behavior to the scalp alone (Figure 3(e)). An increase of 11% at 450 nm up to 39% at 889 nm was obtained by OCS+implant (Figure 3(e)). Maximum optical transmittance of 89% was obtained at 889 nm. Figure 3(f) compares light transmission through OCS+implant and scalp+skull. The bars show the maximum optical transmission of OCS+implant in various wavelength ranges of near-infrared (NIR) (700-900 nm), red (565-700 nm), green (485-565 nm), and blue (450-485 nm). A maximum absolute change of 44.5% in the optical transmittance was obtained using OCS+implant compared to scalp+skull.

**Figure 3.**
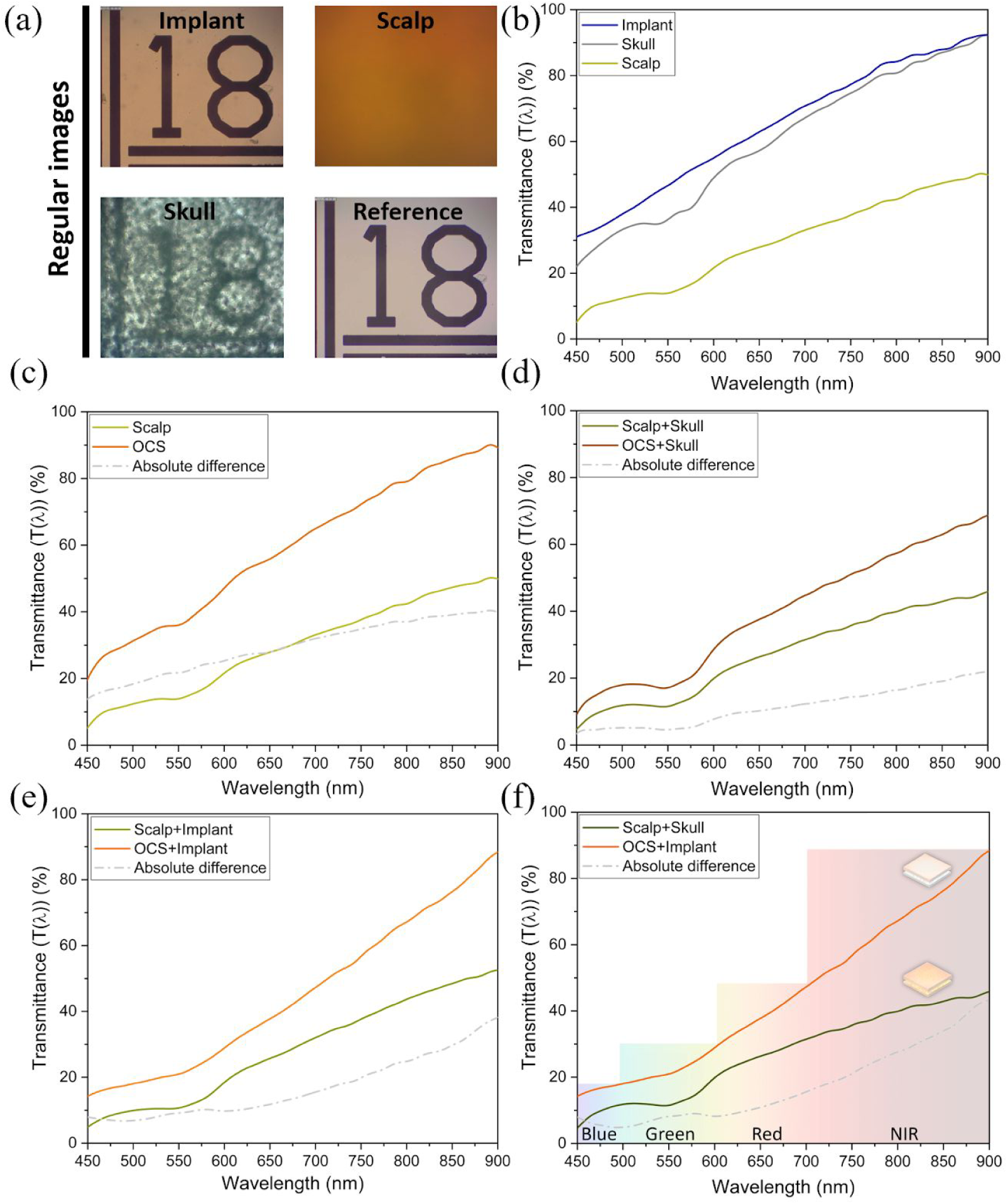
a) Visual comparison of light transmission using photographs of an NBS 1963A resolution target through the WttB implant, skull, and scalp (Experiment 1). The resolutions shown are the 18 cycle/mm target (each black line width is 27.78 μm). b) Optical transmittance of scalp, skull, and implant. c) Comparison of optical transmittance through scalp and OCS. d-f) Comparison of the effect of optical clearing on optical transmittance of the tissue sample stacks; d) scalp+skull vs. OCS+skull, e) scalp+implant vs. OCS+implant, f) scalp+skull vs. OCS+implant.

**Figure 5.**
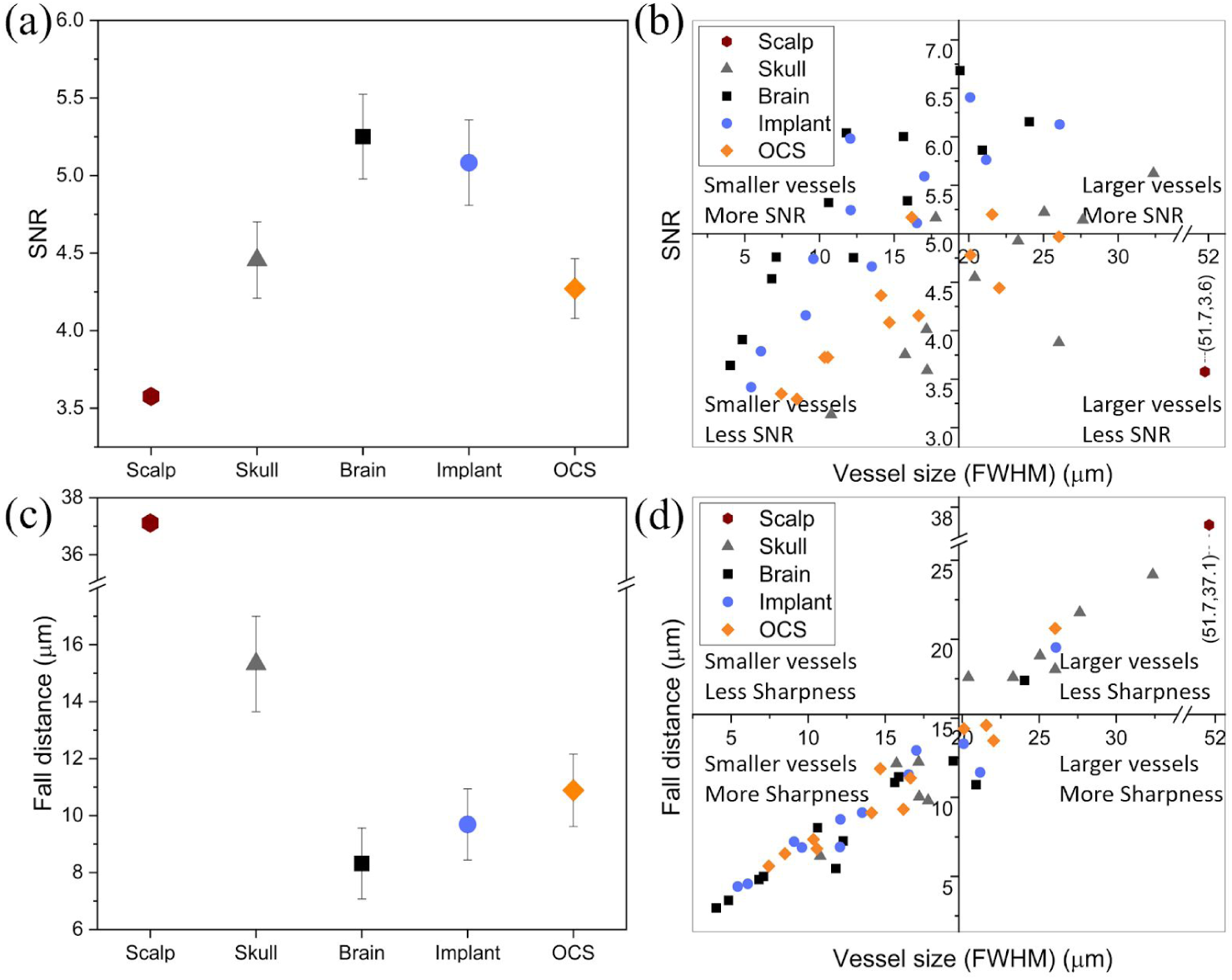
Comparison of LSI image quality over the imaging conditions. a) Mean SNR of contrast intensity along midline profiles on the imaging conditions (Experiment 2). b) SNR vs FWHM for all vessels intersected by line profiles. c) Mean fall distance along the line profiles over the imaging conditions. d) Fall Distance vs FWHM for all vessels intersected. Error bars represent standard error.

### Experiment 2: Brain microcirculation imaging using LSI

The results from Experiment 2 are shown in Figures 4, 5, 6, and 7. As the goal of the WttB implant is to provide optical access to the brain for diagnostic techniques such as microcirculation imaging without scalp removal, we sought to compare LSI through the native scalp with imaging through OCS and implant. An example of LSI images from imaging condition 3 (direct-brain) obtained through different focal planes is shown in Figure 4(a). The image with the highest contrast was chosen for each imaging condition. To make a direct comparison between the imaging conditions, the LSI field of view was kept constant across the conditions. Figure 4(b) shows regular images (showing the LSI field of view) and LSI contrast images at each imaging condition for Mouse 10 (data for Mice 11-12 not shown). It should also be mentioned that blood flow is expected to be altered in response to the invasive cranioplasty surgery (e.g. due to potential reactive hyperemia^52^ (increased blood flow), changes in respiration, etc.). We previously reported the optimal exposure time for our LSI system at different time points after the surgical procedure^24^.

**Figure 4.**
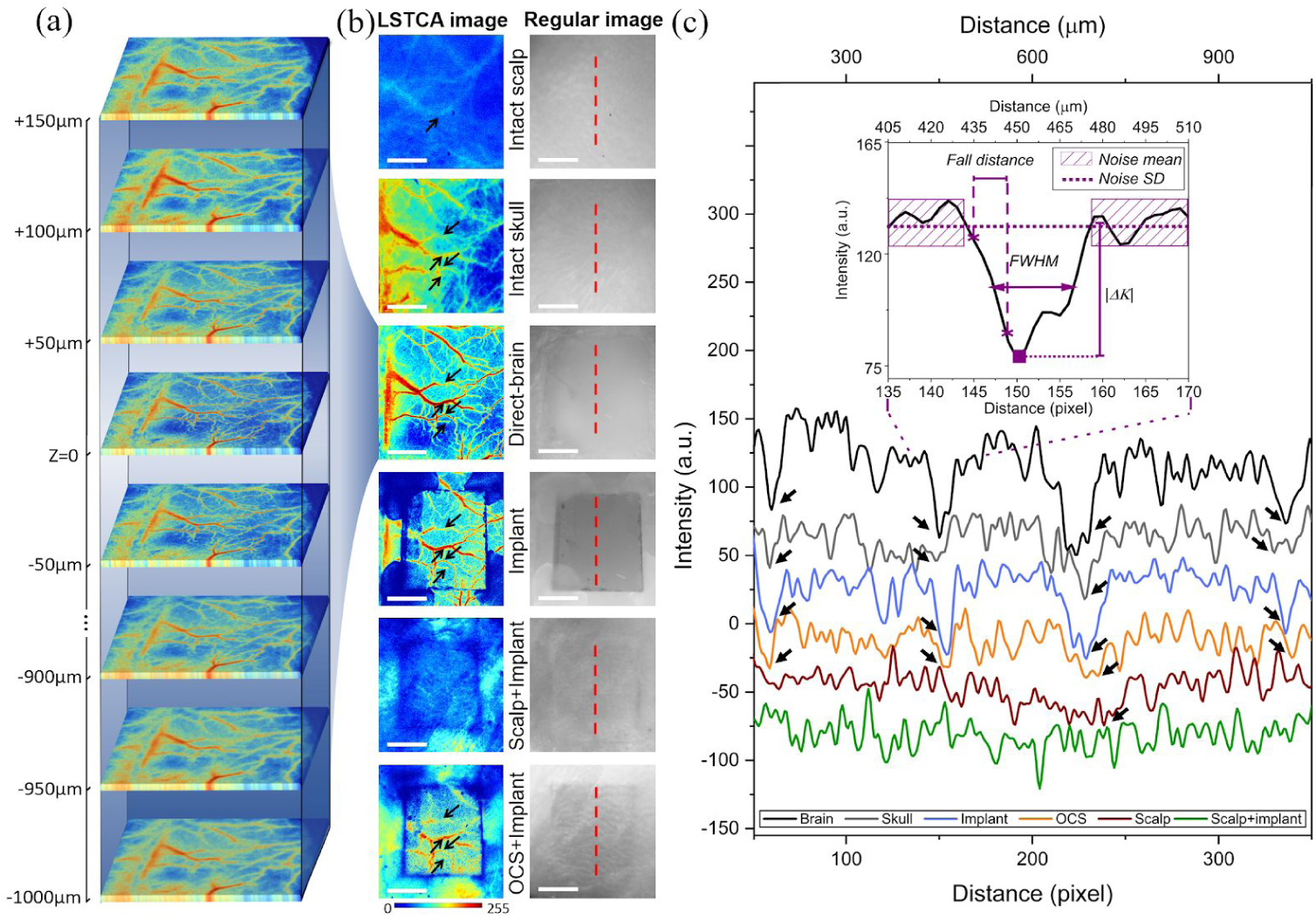
*In vivo* brain microcirculation images over the imaging conditions. a) An example of LSI in imaging condition 3 (direct-brain) focal plane positions (Experiment 2). Focal plane positioned from +150 μm above the surface focal plane (Z = 0) to −1000 μm toward the brain with step size of 50 μm. b) LSI temporal contrast images for 5 imaging conditions of Mouse 10. The right column presents the regular images showing the consistent location along the midline of ROI where line profiles were taken, and the left column shows the corresponding LSI images. c) Example contrast intensity profiles in imaging conditions of 1-6. The arrows in b) and c) show the vessels that are intersected by the midline intensity profiles. Scale bars = 1 mm.

**Figure 6.**
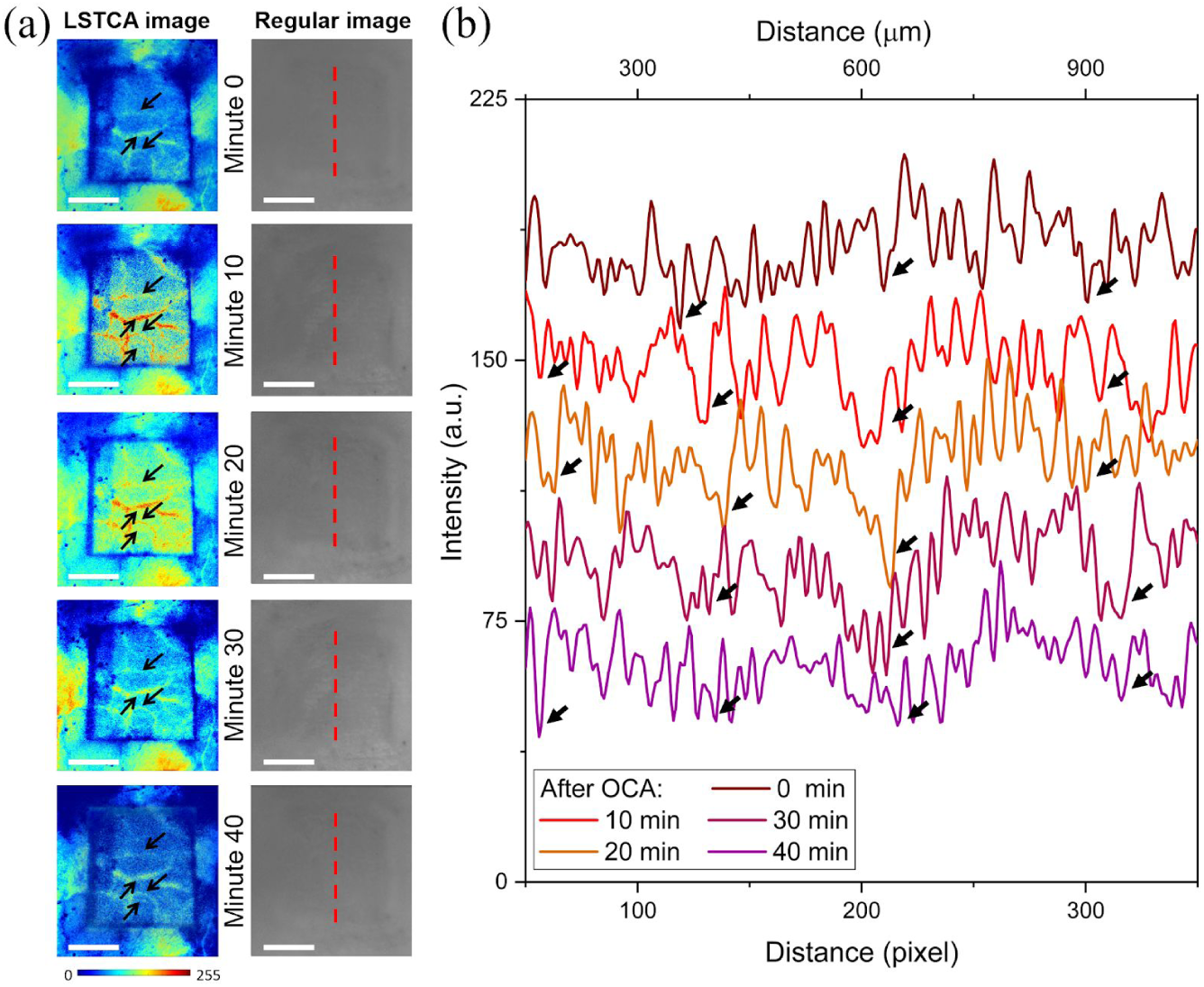
a) *In vivo* brain microcirculation LSI images of OCS+implant (imaging condition 6) immediately after and up to 40 minutes after applying the OCA (Experiment 2). In the right column, the regular images showing the arbitrary locations where line profiles along the midline of ROI were taken, and in the left column, the corresponding LSI images. b) Example contrast intensity profiles in imaging condition 6. Scale bars = 1 mm.

**Figure 7.**
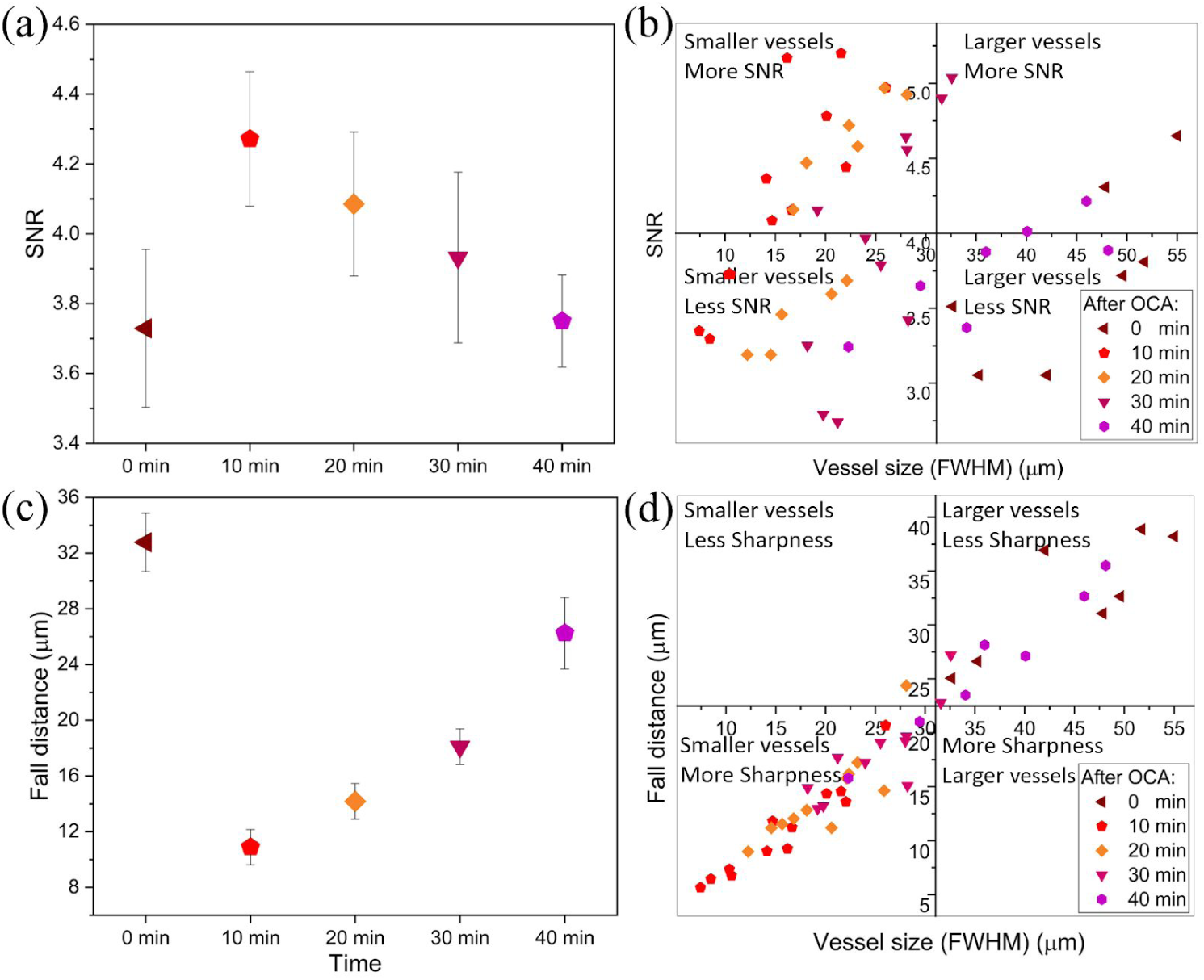
Comparison of LSI image quality of OCS+implant (imaging condition 6). a) Mean SNR of contrast intensity along line profiles on LSI images (Imaging condition 6) immediately after and up to 40 minutes after applying the OCA (Experiment 2). b) SNR vs FWHM for all vessels intersected by line profiles. c) Mean fall distance along the line profiles over the time points. d) Fall Distance vs FWHM for all vessels intersected. Error bars represent standard error.

Line intensity profiles were analyzed for the intact scalp (i.e. condition 1), intact skull (i.e. condition 2), direct-brain (i.e. condition 3), implant (i.e. condition 4), scalp+implant (i.e. condition 5), and OCS+implant (i.e. condition 6) contrast images. To avoid selection bias, the location of these line profiles were chosen arbitrarily at the ROI mid-points, as depicted in the regular image in Figure 4(b). The midlines intersected 3 to 4 vessels like the example shown with arrows in Figure 4(b) and 6(c) for Mouse 10. From these intensity profiles, peak intensity and noise (*ΔK* and *σK*_*n*_) values were determined (as described in the Methods section and illustrated in the inset of Figure 4(c)). Then signal-to-noise ratio (SNR) and fall distance were calculated (see Figure 4(c) for an example trace with the 10–90% fall distance measurement illustrated). Only one brain vessel in Mouse 10 was detectable in the intact scalp LSI image which was used for further comparisons. The scalp+implant did not visualize any vessels which could be considered for comparing SNR, sensitivity, and fall distance.

Data for each imaging condition was averaged between the vessels and the mean SNR for each imaging condition is shown in Figure 5(a). As shown in the Figure, the direct-brain gave the highest SNR, and implant, intact skull, and OCS+implant have lower SNR values respectively; while the intact scalp does not show detectable vessels (only one vessel in Mouse 10). Additionally, the FWHM of each vessel in the line profiles were taken as vessel diameter. Figure 5(b) shows a plot of vessels intersected by the profile line across the imaging conditions, with SNR plotted against FWHM (i.e. vessel diameter). Next, using the 15 LSI temporal contrast images corresponding to these imaging conditions for Mice 10-12, we sought to compare the sharpness (i.e. fall distance) of the vessels in the images to compare the image resolution. Values for each imaging condition were averaged between the vessels and the mean fall distance for each imaging condition is shown in Figure 5(c). A plot of all vessels intersected by the profile line, with absolute fall distance (i.e. vessel sharpness) plotted against FWHM (i.e. vessel diameter) is shown in Figure 5(d).

Considering the temporal effect of OCA on scalp tissue optical transparency (Experiment 1), we sought to quantitatively evaluate this effect on the LSI imaging quality. Figure 6(a) shows regular and LSI images of OCS+implant (imaging condition 6) immediately after and up to 40 minutes after applying the OCA for Mouse 10 (data for Mice 11-12 not shown). The regular images and LSI visually show the temporal change in scalp transparency. Line intensity profiles like the example shown in Figure 4(b and c) were analyzed to determine peak intensity and noise to calculate signal-to-noise ratio (SNR) and fall distance.

Data for each time point (minute 0, 10, 20, 30, and 40) was averaged between the vessels and the mean SNR for each time point after applying OCA (imaging condition 6) is shown in Figure 7(a). The minute-10 contrast image shows the highest SNR (imaging condition 6) which then gradually decreases at minute 20 and 30 and approximately equals to the initial value (minute 0) at minute-40. Figure 7(b) shows a plot of vessels intersected by the profile line across the imaging conditions, with SNR plotted against FWHM (i.e. vessel diameter). Alongside, using the 15 LSI images corresponding to these time points for Mice 10-12, we compared the sharpness (i.e. fall distance) of the vessels in the images to compare the image resolution. Fall distances for each imaging time point were averaged between the vessels and the mean fall distance is shown in Figure 7(c). A plot of all vessels intersected by the profile line, with absolute fall distance (i.e. vessel sharpness) plotted against FWHM (i.e. vessel diameter) is shown in Figure 7(d).

## Discussion

The application of transparent cranial implants holds the transformative potential for facilitating the diagnosis and treatment of a wide variety of brain pathologies and neurological disorders. We have recently reported the properties of our transparent nc-YSZ implant including an excellent ageing resistance evaluated through zirconia-based surgical implants ISO tests^43^. The biocompatibility of zirconia-based implants and the OCA (PG and PEG) have also been reported previously^53,54^. A possible application for our proposed idea is when a craniotomy/craniectomy is needed in a treatment procedure. Then, implementing this implant in the current cranial prostheses or a separate implantation of WttB implant (e.g. on Burr holes) enables delivery to and/or acquisition of light from the brain, in real-time, without the need for scalp removal for various post-operation applications.

In this study, we assessed the light delivery improvement by WttB implant coupled with OCS in a set of *ex vivo* optical characterization measurements. Additionally, we evaluated the feasibility of through-scalp brain microcirculation imaging using the implant *in vivo*. LSI temporal contrast imaging quality (i.e. SNR and sharpness) of cerebral blood vessels imaged through the implant and OCS was calculated.

*Ex vivo* optical transmittance measurements were performed in Experiment 1. The scalp optical clearing results, shown in Figure 2 and Figure 3(c), were in agreement with the previous reports of using PG-PEG agents^48,49,55^. More effective OCAs have been introduced recently and although they have shown a higher transparency, they have not been widely accessible and used^56^. The optimum effect resulted within 10-20 minutes after OCA application. The maximum optical transmission was obtained at minute 11 and wavelength of 890 nm. The solid black lines in Figure 2(a), indicate the zones with specific ranges of optical transparency in which the optimum time and wavelength can be found. OCA provided 13 minutes of transparency in the highest zone (33% - 36% increase in transparency) for wavelengths in the range of 800-900 nm. As our optical window of interest is around wavelength 810 nm for LSI (Experiment 2), the temporal change in that wavelength was extracted from Figure 2(a) and shown in Figure 2(b). A time-window of 13 minutes, from minute 5 to minute 18, associated with the optimum transparency (86%-89%). The effect of OCA on imaging quality through the scalp is visualized in Figure 2(c) and is in agreement with the optical transmission measurements. The highest transmitted light intensity resulted in minute 10. The increased visibility of resolution target through OCS reveals the reduced scattering by the OCA^49^. Since our optical measurements are based on collimated light transmittance we can not directly compare our results to the previous works and our results were used to relatively compare optical transmittance values in the difference conditions. In Figure 3(a), the implant appears as a shade of orange. The implant can be also optimized for various wavelength ranges^57^. The skull tissue transmits the light, although the texture of the cranial bone notably decreases the spatial resolution. The pristine scalp tissue has a very low light transmission. As shown in Figure 3(b), the implant has a higher optical transmittance (up to 12%) than skull. It should also be noted that while the optical transmittance of the implant is an improvement over the skull, the mouse skull is inherently transparent itself,^58^ which is not the case in larger animals or humans. The scalp showed a lower transmittance compared to skull and implant (maximum difference of 39%). This indicates that the scalp is the limiting factor for light transmission within the tissues above the mouse brain. The hemoglobin optical absorbance signature (520-620 nm) is noticeable in scalp tissues (unlike the implant) limiting the access in that wavelength range^59^. Figure 3(f) compares light transmission through OCS+implant and scalp+skull. The bars depict the maximum light transmission through OCS+implant including 89% in NIR range (700-900 nm), 50% in red range (565-700 nm), 24% in green range (485-565 nm), and 20% in blue range (450-485 nm). The enhanced transparency of the YSZ implant coupled with the OCA shows promising features to facilitate various photosensitizer activation-based therapies. Photobiomodulation (600-1064 nm)^1,2,60–62^, PDT(405-900 nm, mainly in red range)^4,63^, optogenetic (400-630 nm)^5,9,64^ are a few examples of optical techniques that could benefit from the improved transmission in offered by the YSZ implant. Complementarily, any potential adhesion of biochemical agents and/or tissue growth on the implant (e.g., fibrotic tissue, proteins, cell adhesion) could be potentially monitored over time^65^.

In Experiment 2 we performed *in vivo* brain microciculation imaging. Laser speckle imaging has become a useful tool for brain blood flow applications as the images it produces contain functional information (i.e. relative blood velocity) in addition to showing structure of the vessel networks^13,66,67^. As we previously reported, exposure time of 6 ms gave the highest SNR for imaging through WttB implant using our optical setup^24^. SNR and sharpness (fall distance) of vessels are evaluated for an identical field of view between the various imaging conditions, allowing us to directly compare the SNR values of intact scalp (i.e. condition 1), intact skull (i.e. condition 2), implant (i.e. condition 4), scalp+implant (i.e. condition 5), and OCS+implant (i.e. condition 6) to the reference image (direct-brain, condition 3) to evaluate the sensitivity. A visual comparison over the imaging conditions in Figure 4(b) shows the brain microvessels were not detected through the intact scalp and only a blurred vision of the largest vessel in Mouse 10 can be seen and hardly found in the profile line (Figure 4(b)). More details are visible in the intact skull image, and likewise, the brain vessels can be seen and detected through the profile. The direct-brain image has the highest contrast and resolution, and the vessel can be clearly seen in the intensity profile. Likewise, vessels are observed in the implant image, however, a lower contrast intensity is noticeable compared to the direct-brain image. The contrast of the vessels to the background (brain tissue) and resolution (sharpness) is notably higher compared to skull image. In OCS+implant image, brain vessels and microvessel are visible and detectable in the intensity profile. Compared to intact scalp image, significantly higher contrast and resolution are observed. However, the contrast and resolution are lower than those of the direct-brain and implant images. The uncleared scalp+implant did not visualize vessels.

Quantitatively, the direct-brain image with average SNR of 5.25 had the highest signal (contrast) to noise ratio. As the size of vessels and velocity of blood flow is not expected to differ on average between the imaging conditions, the apparent increase in SNR and vessel diameter imaged through the scalp, skull, implant, and OCS+implant is likely due to the blurring of the image. OCS+implant showed a higher SNR of 4.25 compared to intact scalp with SNR of 3.58. Implant and intact skull images had SNR values of 5.08 and 4.45, respectively. OCS+implant imaging resulted in Sensitivity of 81%, while intact scalp showed Sensitivity of 68% (only one detected vessel was considered). Implant and intact skull imaging sensitivities were 96% and 84%, respectively. As seen in Figure 5(b) the only visible vessel in the intact scalp image appeared very large (FWHM = 71.7 μm) and also had a SNR of 3.6 while in the implant+OCS image, the vessels were all detected and are visible with vessel sizes of 7.5 μm to 26 μm. In Figure 5(b), implant image vessels showed a similar pattern to the direct-brain vessels although they are slightly shifted toward lower SNR and higher size.

The intact scalp image resulted in the lowest averaged sharpness (the highest fall distance of 37.1 μm) while OCS+implant showed fall distance of 11μm (see Figure 5(c)). The implant images had a lower fall distance of 9.6 μm compared to intact skull image with fall distance of 15.3 μm. The skull and OCS scattering both disorder the speckle pattern that was created by the brain hemodynamics. The reduction in border sharpness of the vessels imaged through the intact skull vs OCS+implant suggests that the skull has a higher blurring effect as seen in Figure 4(b) and Figure 5(c). The direct-brain images with fall distance of 8.2 μm showed the highest vessel sharpness. In Figure 5(d), the data from OSC+implant, implant, and direct-brain fell into a similar cluster placed mostly in the lower left quadrant associated with lower vessel size and higher sharpness. On the other hand, a half of intact skull vessels were detected in the high vessel size and low sharpness quadrant. The intact scalp image shows the lowest sharpness with fall distance of 37.1 μm. The SNR and sharpness of the OCS+implant images showed an improvement over the intact scalp image. We believe this improvement will be significant in the case of larger animals or humans in which the thicker skull bone totally inhibits any optical-based through-skull cerebral blood flow visualization.

In Figure 7(a), SNR increases from minute 0 to minute 10 and then gradually decreases until minute 40, which is in agreement with Experiment 1 (Figure 2(b,c)). The trend of the OCA temporal effect is also in agreement with previous studies on LSI and OCAs^68,69^. A relatively high standard deviation of the data (low certainty and more similarity between the averaged values) can be noticed in Figure 7(a). In Figure 7(b), the vessels at minutes 10, 20, and 30 fell in the smaller vessel size quadrants (left half) covering both lower and higher SNR quadrants. On the other hand, the vessel at minutes 0 and 40 mostly fell in the lower SNR quadrants (lower half) including the smaller and larger vessel size quadrants. This suggests that within the time points with higher effect of OCA (minutes 10, 20, 30), the effect on detected vessel size is more obvious than the effect on the SNR value. The effect of OCA on the imaged vessel sharpness (Figure 7(c)) shows relatively lower deviation over the time points compared to that of the SNR. Additionally, the separated clusters of imaged vessels at minutes 10 and 20 in Figure 7(d) fell in the lower size and higher sharpness suggests that the OCA effect is more noticeable in sharpness (resolution) of the image than SNR.

We compared the results of Experiment 2 obtained through intact skull and OCS+implant showing OCA was more effective in reducing the blurriness (i.e. the reduced size and increased sharpness of the detected vessels) while it did not increase the SNR values of OCS+implant imaged vessels compared to that of intact skull. Then we considered the results of Experiment 1 which show skull also provided higher optical transmittance than OCA+implant (Figure 3). Experiment 1 suggests that the average amount of incoherent light transmitted through skull was higher than that of the OCS+implant and consistently, Experiment 2 depicts skull provided a lower SNR for LSI. The increase in the sharpness and decrease in the vessel size are due to the reduced scattering in the scalp by OCA^69^ while it did not provide higher light transmission and SNR (in LSI) which rely on both scattering and absorption. This shows OCA was more effective in reducing the scattering than the absorption in scalp. Although the OCA’s dominant effect on scattering has been widely reported, there are two points regarding the effect of OCA on SNR (in LSI) that should be added here. First, the reduced scattering and increased transparency by OCA resulted in the ability to visualize more scalp textural details (see Figure 2(c)), which is considered as noise in the SNR calculation and decreases the final value. Second, OCA creates dynamic spatiotemporal behavior in the optical properties such as index of refraction which consequently affects the optical paths of transmitted laser light through the tissue. This might result from the penetration of OCA and water in the tissue after applying the OCA. This effect (which we call “dynamic transparency”) affects coherence-based imaging modalities with long data acquisition time such as LSI (with camera exposure time of ∼5-25 ms) by decreasing the contrast between dynamic and static region resulting in lower signal and SNR values.

Microcirculation plays a critical role in physiological processes such as tissue oxygenation and nutritional exchange^11,70^. Monitoring the spatiotemporal characteristics of microcirculation is crucial for studying the normal and pathophysiologic conditions of tissue metabolism. For example, microcirculation monitoring is useful for assessing microcirculatory dysfunction due to disease conditions such as type 2 diabetes, peripheral vascular disease (PVD), atherosclerotic coronary artery disease, obesity, heart failure, Alzheimer’s, schizophrenia and hypertension, among others^71–73^. In addition, quantification of dynamic blood flow, angiogenesis, and vessel density are critical for monitoring the progression of wound healing^74^. Although high-resolution vascular network mapping is possible using imaging modalities such as computed tomography (CT), these approaches require injection of contrast agents and pose disadvantages such as radiation exposure. Existing non-invasive methodologies (including LSI through skull) are inadequate to study brain blood flow at microvessel resolution^75^. Optical accesses such as OCS coupled with WttB implant are thus important tools for research and can become important enablers of clinical diagnostics and therapy involving cerebral microvessels. Figure 8 visually illustrates the enhancement in optically accessing the brain hemodynamics after the implantation and through closed scalp (OCS+implant). An example region in the flowmetric image shown with dashed line was compared to direct-brain and intact scalp images. The vessel intersected by the black line (V1) has a width of 22.5 μm and the vessel intersected by the purple line (V2) has a width of 16.8 μm. Precise velocity information, particularly of microvessels, appeared through the OCS+implant, while the inhibited signal through intact scalp and skull obscures flow determinations.

**Figure 8.**
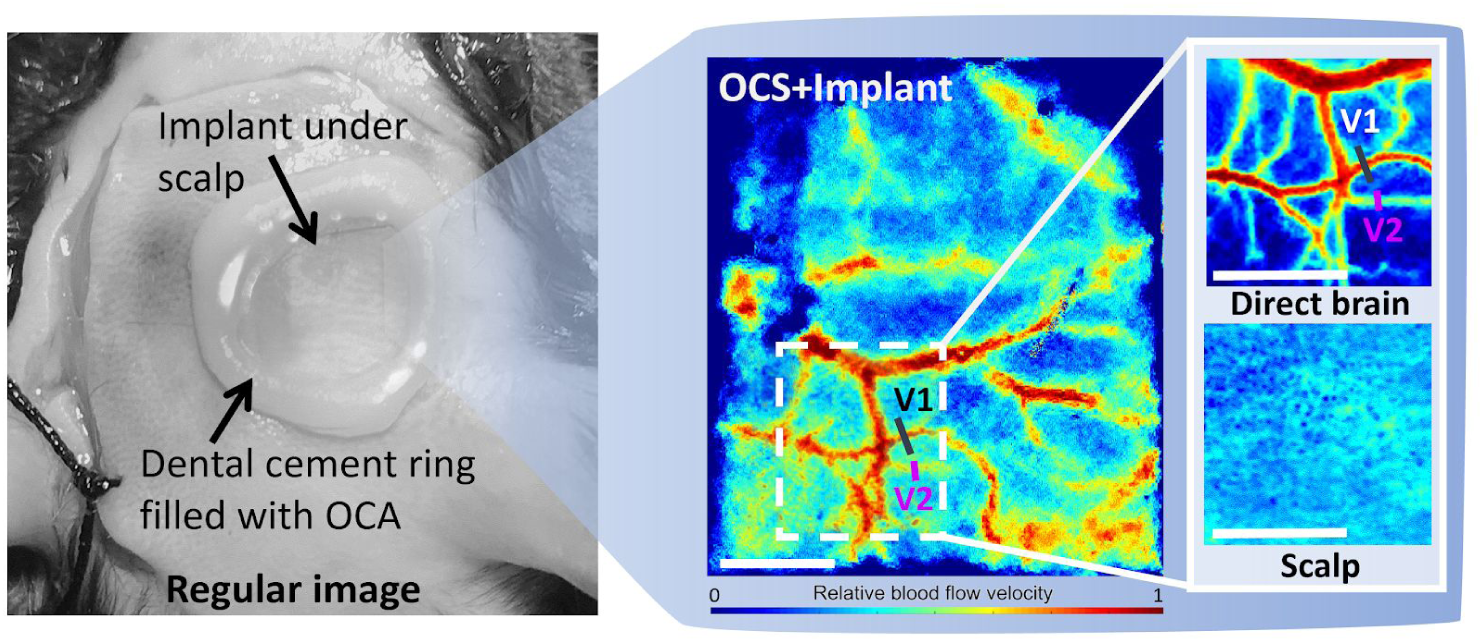
A regular image of Mouse 10 head showing the provided brain flowmetric imaged obtain through scalp (Experiment 2). The implanted WttB prosthesis and a drop of OCA on the closed scalp was able to provide access to the brain hemodynamic (shown in the inset). An example region in the flowmetric image (shown with dashed line) was compared to the reference (direct-brain) and intact uncleared scalp images. The vessel intersected by the black line (V1) has a width of 22.5 μm and the vessel intersected by the purple line (V2) has a width of 16.8 μm. Scale bar = 500 μm.

Creating novel windows for brain studies has been gaining attention recently^35,76–78^. Some of these studies, involving OCAs applied to the scalp overlying native skull, have shown limited success due to optical losses and scattering in the skull^32^. Future studies by our group will explore this combined OCA-WttB strategy for chronic imaging in awake and behaving animals through closed scalp, to study cerebrovascular hemodynamic responses to a photoactivation stimulation. There are several limitations to the current study. The sample sizes used in Experiments 1 and 2 were small (n = 3), and further experiments are needed to confirm the reproducibility of these findings. While a permanent cranial implant can allow for less invasive imaging of the brain at later time points, it requires an initial implantation surgery which carries associated risks such as infection.

In this study, optical access to mouse cerebral microvasculature was achieved using a transparent cranial implant and optical clearing of the scalp. Transmittance peaked at 89% around 10 minutes after application of the optical clearing agents, and slowly decayed, allowing a window of ∼13 minutes suitable for laser speckle imaging. Strategies for providing ongoing optical access to the brain without repeated craniectomy and/or scalp retraction, like the one presented here, could one day improve clinical imaging and optical diagnosis and treatment of brain disease.

## METHODS

### Transparent stabilized-zirconia cranial implants

Transparent nanocrystalline 8 mol% YO_1.5_ yttria-stabilized zirconia (nc-YSZ) implant samples were produced from a precursor YSZ nanopowder (Tosoh USA, Inc., Grove City, OH, USA) densified into a transparent bulk ceramic via Current-Activated Pressure-Assisted Densification (CAPAD) as described previously.^79^ The thickness of the resulting densified YSZ discs were reduced from 1 mm to ∼440 μm by polishing with 30 μm diamond slurry on an automatic polisher (Pace Technologies, Tucson, Arizona, USA). The samples were then polished with successively finer diamond and silica slurries ranging from 6 to 0.2 μm. Samples were sectioned into rectangles of approximately 2.1 00D7; 2.2 mm using a diamond lapping saw (WEIYI DTQ-5, Qingdao, China), followed by sonication in acetone and thorough rinsing in water. Optical, mechanical, and ageing properties of the material were reported previously^38,57,79^.

### Animals

C57BL/J6 WT (#000664) mice were obtained from Jackson Laboratories. Animals were maintained under a 12-hour light/dark cycles and were provided food and water *ad libitum*. All experiments were performed with approval from the University of California Animal Care and Use Committee and in accordance with the National Institute of Health Animal Care and Use Guidelines. Males between 8 – 12 weeks of age were used in this study.

### Preoperative surgical preparation and anesthesia

Mice were anesthetized with isoflurane inhalation (0.2-0.5%) and given an i.p. injection of ketamine and xylazine (K/X) (80/10 mg/kg). Mice were aseptically prepared for surgery and secured in a stereotaxic apparatus. Artificial tear ointment was applied to the eyes to prevent drying. Toe pinch reflex was used to measure anesthetic depth throughout the experiment, and supplemental doses of K/X were administered as needed.

### Optical clearing agent

A mixture of two biocompatible chemicals, PEG-400 (PEG) and Propylene Glycol (PG) (Fisher Scientific, California, US), were used as scattering reducer and penetration enhancer, respectively, at a volume ratio of 9:1 at room temperature.^53,56^

### Experiment 1: Ex vivo optical transmittance measurements

#### Scalp and skull tissue acquisition and preparation

A sagittal incision was made to the left of the midline, and a square section of the scalp was excised. Periosteum was removed from the skull, and a craniectomy was performed with a surgical drill and carbide burr to excise a square section of skull over the right parietal lobe, with dimensions slightly larger than the implant (2.6 x 2.6 mm) (see Figure 9(a)). Excised full thickness scalp and cranial bone from above the right parietal lobe were rinsed briefly in saline solution to remove excess blood. The thicknesses of the samples were measured to be: 440 ± 1μm, 159 ± 1μm and 710 ± 20μm for the YSZ implant, the mice skull and scalp, respectively. Hence, the YSZ implant is 2.5 times thicker than the mouse skull. Following excision of tissue, mice were euthanized by intraperitoneal (I.P.) injection of pentobarbital solution.

**Figure 9.**
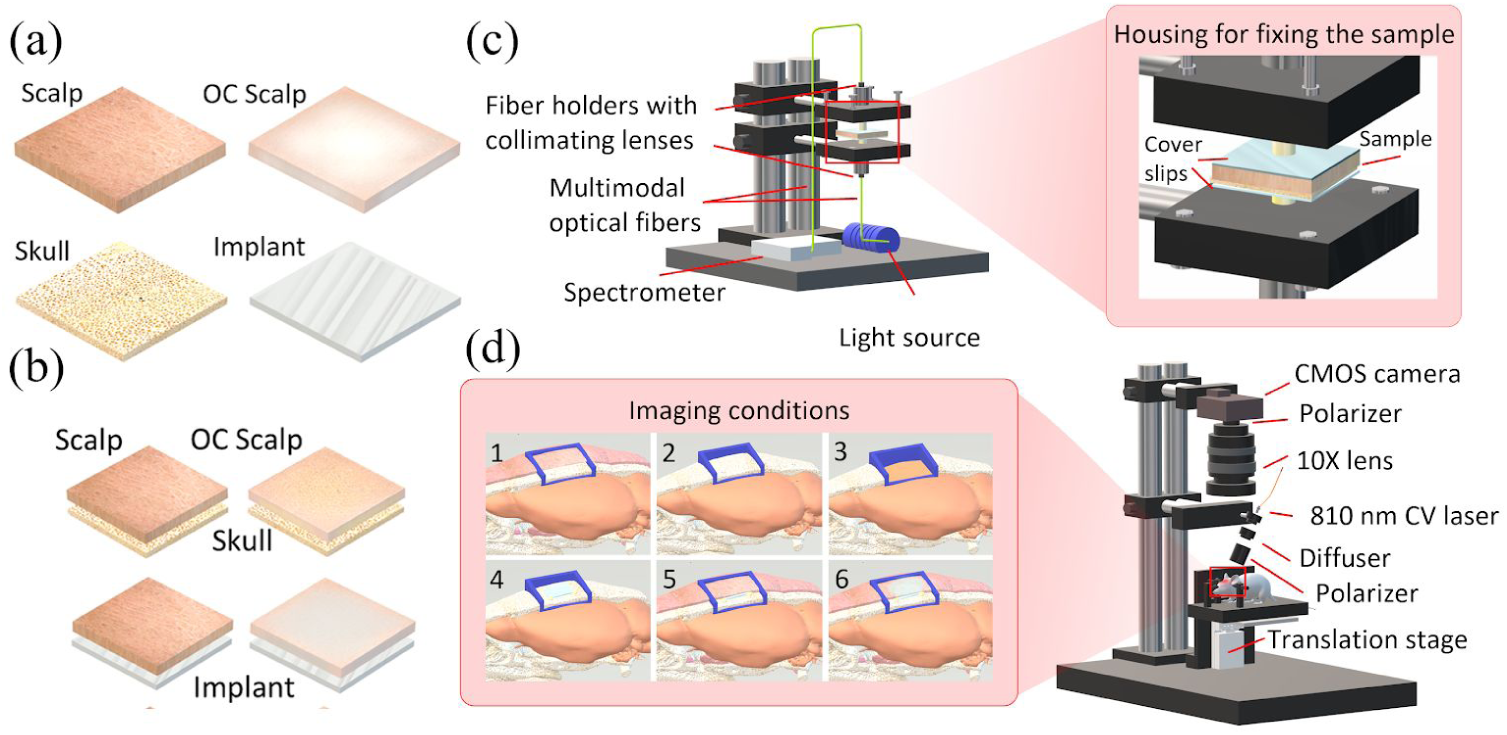
Optical characterization and LSI experimental setups. a,b) Samples used in the *ex vivo* optical characterization (Experiment 1). a) Single layer samples of implant, skull, scalp, and OCS. b) Stacked sample arrangement used to compare the spectral transmittance between: first row) scalp and OCS placed on skull, second row) scalp and OCS on implant. c) *ex vivo* experimental setup for optical transmittance measurement (Experiment 1). d) *in vivo* LSI setup (Experiment 2). The inset shows a schematic of the imaging conditions 1-6, with the blue tetragonals representing the imaging fields of view on murine cranium.

The OCA was prepared and topically applied at room temperature. A thin layer of the OCA was applied on the scalp sample and remained for 50 minutes.^53,56^ Characterization measurements were performed before and immediately after applying the OCA, and the change in transmittance for each sample was monitored every minute over the 50-minute period.

#### Optical transmittance measurement setup

Optical transmittance measurements of the different samples used in this study were obtained through optical spectrometry. The setup used to obtain the transmittance spectra incorporates two multimode optical fibers (P400-2-VIS-NIR, Ocean Optics, FL) attached to individual fiber holders including visible-NIR collimating lenses (MP-74-UV, Ocean Optics, FL, with focal length f = 10mm, lens diameter D = 5mm and N A = D/2 f = 0.4). As depicted in Figure 9(c), the holders were attached to a mechanical rail allowing for adjustment of the distance between the fibers and to incorporate a sample holder. After the sample was secured within the holder, the fiber holders were tightly joined together to mitigate detrimental effects from ambient light and back reflections. The light source used for these measurements was a visible-NIR source (HL2000 FHSA, Ocean Optics, FL) launched into one of the optical fibers. The beam exiting the launching fiber then traverses the sample and is collected by the other fiber, which is connected to a solid-state spectrometer (SD2000, Ocean Optics, FL) to obtain the optical transmission spectra. Spectra were acquired averaging 10 measurements, with an integration time of 300 ms, in the 450-900 nm wavelength range. For all the measurements, the collimated transmittance (T(λ)) was calculated as the ratio of light transmitted through the sample to the total incident light, Equation 1:

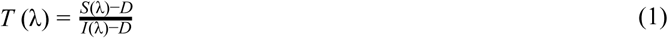

where λ is the wavelength, S is the measured spectral intensity, I is the total light incident and D represents the reference reading under dark conditions (i.e., no light impinging on the sample). Δ*T* (λ) is absolute change in transmittance, Equation 2.

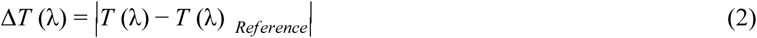

A set of photographs of a resolution target imaged through a scalp tissue was also captured before and up to 40 minutes after applying the OCA using a metallurgical microscope (MT 8530, Meiji Techno, Japan).

#### Experimental design

For Experiment 1, the procedure used for measuring the ballistic transmittance through the different tissues and the implant was similar to procedures reported previously for spectroscopic measurements on soft tissue^80–82^. Three sessions of measurements were performed. In each session, optical measurements were repeated n=3 times using freshly excited samples from 3 new mice. All of the samples were placed between two glass microscope coverslips to obtain the transmittance spectra. In the first session, scalp and skull samples were excised (from Mice 1-3) and optical transmittance of single layer samples were measured (Figure 9(a)). After obtaining implant, skull, and scalp spectra, the OCA was applied on scalp and the spectral transmittance of OCS was acquired over time (Figure 9(a)). Stacking arrangements were then used for the second and third sessions of measurements allowing for evaluating the effects of each layer on the spectral features of the sample. This further allows for comparing the spectral features of the skull and the YSZ implant under similar conditions. In the second session, 3 new scalp and skull samples were excised (from Mice 4-6) and spectra were obtained for the stacked arrays of the scalp placed on top of the skull (Figure 9(b), upper row). Then, the OCA was applied on the scalp and samples (OCS-on-skull) were measured over time. In the third session of measurements, 3 new scalp samples were excised (from Mice 7-9) and the stacked array of the scalp placed on top of the implant was measured (Figure 9(b), lower row). Then, the OCA was applied on the scalp and samples (OCS-on-implant) were measured over time.

### Experiment 2: In vivo brain hemodynamics imaging

#### Mouse model preparation

The mouse model for this study was designed to provide optical access to the right parietal lobe of the brain through the OCS and WttB implant (Figure 10(a)). A sagittal incision was made to the left of the midline, and the scalp retracted to expose the skull (Figure 10(b)). Periosteum was removed from the skull, and a craniectomy was performed with a surgical drill and carbide burr to remove a square section of skull over the right parietal lobe, with dimensions slightly larger than the implant (Figure 10(c)). The YSZ implant was placed within the craniectomy directly on the intact *dura mater*, and dental cement was applied to each of the four corners of the implant to prevent displacement (Figure 10(d)). Dental cement was cured with blue light exposure for 20 seconds. Imaging with LSI was conducted before and after opening the scalp (imaging conditions 1 and 2, respectively), immediately after the craniectomy procedure, while the scalp was still open (imaging condition 3), and after the implantation (imaging condition 4). Then, the retracted scalp was placed back over the implant and a ring-shape barrier (for holding OCA) was made on top of the scalp using dental cement and cured with blue light (Figure 10(e)). LSI imaging was conducted before and immediately after adding a drop of OCA inside the dental cement holder (imaging condition 5 and 6, respectively). The OCA were applied after the implantation and replacement of the scalp on the head (Figure 10(e)). The solid ring of dental cement placed on the closed scalp surrounded the OCA and inhibited the agents from leaking out of this region of interest. A drop of OCA was carefully added inside the ring and imaging was performed before and immediately after applying the OCA.

**Figure 10.**
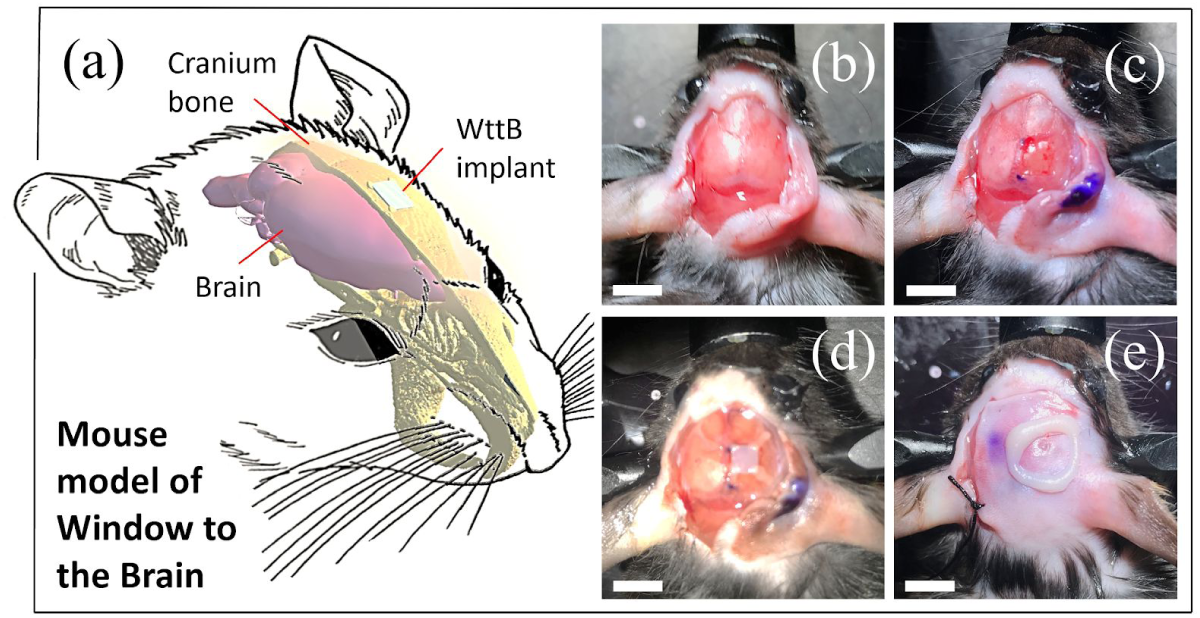
a) A schematic of the mouse model used for this study (Experiment 2) showing the location of optical access. b-e) Surgical preparations (scale bars = 4 mm), retracted scalp to expose the skull (b), craniotomy and implantation (c,d), and replacing the scalp and making a ring-shaped OCA holder (e).

#### Laser speckle imaging setup

For LSI, an 810 nm continuous wave NIR laser (DL808-7W0-O, CrystaLaser, Reno, NV, USA) was used to illuminate the region of interest with an incident power of 100 mW at a 45° incidence. While most LSI studies use visible wavelengths for illumination, we chose 810 nm to reduce reflectance and increase transmittance through the WttB implant.^24^ The 810 nm laser was expanded using a pair of negative-positive lenses (KPC043, −25 mm EFL and KPX094, 100 mm EFL, Newport, Irvine, CA, USA) and speckles were generated and homogenized using an engineered diffuser (ED1-C20-MD, Thorlabs, Newton, NJ, USA). Diffused laser light was shown onto the intact scalp (i.e. condition 1), onto the intact skull (i.e. condition 2), onto the dura mater and cortex after craniectomy (i.e. condition 3), through the WttB implant (i.e. condition 4), onto the closed scalp (i.e. condition 5), and onto the closed scalp after applying the OCA (i.e. condition 6). The crossed polarized (LPVIS100, Thorlabs, Newton, NJ, USA) reflected light from the illuminated region was captured by a 12-bit CMOS camera (DCC1545M, Thorlabs, Newton, NJ, USA) equipped with a 10X zoom microscope (MLH-10X, 152.4 mm WD, Computar, Torrance, CA, USA). The aperture and magnification of the zoom microscope were carefully chosen to ensure that the speckle size at the image plane was approximate to the area of two pixels in the CMOS chip. A schematic of the imaging system is shown in Figure 9(d).

To ensure the LSI system was focused on the correct plane, for each imaging condition, the focal plane was shifted between positions (from +150 μm above the surface focal plane (Z = 0) to −1000 μm toward the brain) with step size of 50 μm using a motorized translation stage (MTS50E, Thorlabs, Newton, NJ, USA). At each position, a sequence of 100 laser speckle images were captured at exposure time of 6ms (per our previous report on optimized LSI exposure time)^24^ at a speed of 14 frames per second. Then, the laser speckle images were analyzed to obtain the contrast image.

#### Image processing and data analysis

The contrast-resolved LSI images were constructed based on temporal statistical analysis of laser speckle which has been proven to preserve spatial resolution.^58^ Experimental results have indicated that temporal speckle contrast analysis could expressively suppress the effect of the static laser speckle pattern formed by the stationary superficial reflection and scattering tissue on the visualization of blood flow^58,68,83–85^. Suppressing this effect makes temporal contrast analysis an ideal method for imaging cerebral blood flow, and in this study we assess whether this method can resolve blood flow across the skull, scalp, OCS, and implant. The temporal contrast, *K*_*t*_, of each image pixel in the time sequence was calculated using Equation (3),^83^

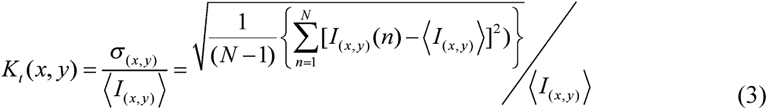

where *I*_*x,y*_*(n)* is the CCD counts at pixel (x,y) in the nth image, N is the number of images acquired, and 〈*I*_*x,y*_〉 is the mean value of CCD counts at pixel (x,y) over the N images. The raw images were process to obtain the contrast image. The contrast images across the height positions were compared to find the correctly focused plane (associated with the highest contrast) for each imaging condition.

We assessed the quality of the speckle contrast images over time in terms of signal to noise ratio (SNR) and vessel sharpness. To quantify signal to noise ratio for each imaging condition, the contrast intensity profile along a vertical line (across the blood vessels) was considered. To avoid selection bias, the location of these line profiles were chosen arbitrarily at the region of interest (ROI) mid-points. The midlines intersected 3 to 4 vessels, and remained the same between the imaging conditions. Equation (4) shows how SNR values were calculated for each imaging condition,

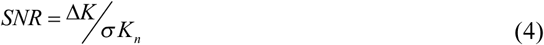

where *ΔK* is the depth of the vessel peak from the baseline (mean noise) and *σK*_*n*_ is the standard deviation of the noise. The SNR values were averaged over the mice (10, 11, and 12) and standard errors were calculated. Sensitivity, which is considered as the ratio of the mean SNR to the mean SNR of the reference, was calculated using Equation (4).

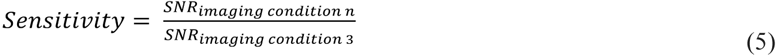

As an indicator of resolution, we compared the sharpness of the vessel edges in images by calculating fall distance (the number of pixels multiplied by the pixel size (∼3 μm)) of the edge of the vessel to go from 10% to 90% of *ΔK* value.^86^ A shorter fall distance corresponds to greater sharpness. The same sampled contrast intensity profile lines were considered for the fall distance calculation. To consider the vessel sizes while comparing SNR and fall distance, full-width half-max (FWHM) of the vessels in profiles were taken as the vessel diameter.^86^

#### Experimental design

After the *ex vivo* optical transmittance measurements in Experiment 1, a session of *in vivo* brain imaging using LSI was performed on 3 new mice (referred to hereafter as Mice 10-12). *In vivo* LSI through the six imaging conditions were performed in succession in each of the mice (n=3). Figure 9(d) inset shows the imaging conditions of intact scalp (i.e. condition 1), intact skull (i.e. condition 2), direct-brain (i.e. condition 3), implant (i.e. condition 4), scalp+implant (i.e. condition 5), and OCS+implant (i.e. condition 6).

## Data availability

The datasets generated during the current study are available from the corresponding author on reasonable request.

## Acknowledgments

National Science Foundation (NSF-PIRE, 1545852 & NSF-EAGER, 1547014), “Beca Mixta” from National Council of Science and Technology of Mexico (CONACYT) (741249), and DGAPA-PAPIIT (Grant IG100519). The authors would like to acknowledge Gottlieb Uahengo for fabricating the YSZ samples.

## Authors Contributions

Study designed: N. D., M. S. C., and G. A. Optical characterization system designed and performed: M. S. C., N. D., and J. A. H. Surgical procedure designed: D. K. B. and C. R. J. Surgical procedure performed: D. L. H and C. R. J. Spectroscopy data analyzed: M. S. C. Image processed and analyzed: N.D. Wrote the manuscript: N.D. and D. L. H. All authors have reviewed the manuscript and agreed regarding its submission.

## Additional Information

The authors declare that there are no competing financial interests.

## References

1. Yun, S. H. & Kwok, S. J. J. Light in diagnosis, therapy and surgery. Nat Biomed Eng 1, (2017).

2. Chung, H. et al. The nuts and bolts of low-level laser (light) therapy. Ann. Biomed. Eng. 40, 516–533 (2012).

3. Naeser, M. A. et al. Significant improvements in cognitive performance post-transcranial, red/near-infrared light-emitting diode treatments in chronic, mild traumatic brain injury: open-protocol study. J. Neurotrauma 31, 1008–1017 (2014).

4. Agostinis, P. et al. Photodynamic therapy of cancer: an update. CA Cancer J. Clin. 61, 250–281 (2011).

5. Gradinaru, V., Mogri, M., Thompson, K. R., Henderson, J. M. & Deisseroth, K. Optical deconstruction of parkinsonian neural circuitry. Science 324, 354–359 (2009).

6. Creed, M., Pascoli, V. J. & Lüscher, C. Addiction therapy. Refining deep brain stimulation to emulate optogenetic treatment of synaptic pathology. Science 347, 659–664 (2015).

7. Ramirez, S. et al. Activating positive memory engrams suppresses depression-like behaviour. Nature 522, 335–339 (2015).

8. Iyer, S. M. et al. Virally mediated optogenetic excitation and inhibition of pain in freely moving nontransgenic mice. Nat. Biotechnol. 32, 274–278 (2014).

9. Bruegmann, T. et al. Optogenetic control of contractile function in skeletal muscle. Nat. Commun. 6, 7153 (2015).

10. Gutruf, P. & Rogers, J. A. Implantable, wireless device platforms for neuroscience research. Curr. Opin. Neurobiol. 50, 42–49 (2018).

11. Schwartz, W. J. et al. Metabolic mapping of functional activity in the hypothalamo-neurohypophysial system of the rat. Science 205, 723–725 (1979).

12. Gusnard, D. A., Raichle, M. E. & Raichle, M. E. Searching for a baseline: functional imaging and the resting human brain. Nat. Rev. Neurosci. 2, 685–694 (2001).

13. Towle, E. L., Richards, L. M., Kazmi, S. M. S., Fox, D. J. & Dunn, A. K. Comparison of indocyanine green angiography and laser speckle contrast imaging for the assessment of vasculature perfusion. Neurosurgery 71, 1023–30; discussion 1030–1 (2012).

14. Scerrati, A. et al. Indocyanine green video-angiography in neurosurgery: a glance beyond vascular applications. Clin. Neurol. Neurosurg. 124, 106–113 (2014).

15. Jonathan, E., Enfield, J. & Leahy, M. J. Correlation mapping method for generating microcirculation morphology from optical coherence tomography (OCT) intensity images. J. Biophotonics 4, 583–587 (2011).

16. Hu, S., Maslov, K., Tsytsarev, V. & Wang, L. V. Functional transcranial brain imaging by optical-resolution photoacoustic microscopy. J. Biomed. Opt. 14, 040503 (2009).

17. Leahy, M. J. Microcirculation Imaging. (John Wiley & Sons, 2012).

18. Devor, A. et al. Frontiers in optical imaging of cerebral blood flow and metabolism. J. Cereb. Blood Flow Metab. 32, 1259–1276 (2012).

19. Tsytsarev, V., Rao, B., Maslov, K. I., Li, L. & Wang, L. V. Photoacoustic and optical coherence tomography of epilepsy with high temporal and spatial resolution and dual optical contrasts. J. Neurosci. Methods 216, 142–145 (2013).

20. Gottschalk, S., Fehm, T. F., Deán-Ben, X. L., Tsytsarev, V. & Razansky, D. Correlation between volumetric oxygenation responses and electrophysiology identifies deep thalamocortical activity during epileptic seizures. Neurophotonics 4, 011007 (2017).

21. Zhang, Q. et al. Wide-field optical coherence tomography based microangiography for retinal imaging. Sci. Rep. 6, 22017 (2016).

22. Klein, K. U. et al. Measurement of cortical microcirculation during intracranial aneurysm surgery by combined laser-Doppler flowmetry and photospectrometry. Neurosurgery 69, 391–398 (2011).

23. Miao, P. et al. Chronic wide-field imaging of brain hemodynamics in behaving animals. Biomed. Opt. Express 8, 436–445 (2017).

24. Davoodzadeh, N. et al. Evaluation of a transparent cranial implant as a permanent window for cerebral blood flow imaging. Biomed. Opt. Express, BOE 9, 4879–4892 (2018).

25. Davoodzadeh, N. et al. Laser speckle imaging of brain blood flow through a transparent nanocrystalline yttria-stabilized-zirconia cranial implant. in Dynamics and Fluctuations in Biomedical Photonics XV 10493, 1049303 (International Society for Optics and Photonics, 2018).

26. Davoodzadeh, N. et al. Optical Access to Arteriovenous Cerebral Microcirculation Through a Transparent Cranial Implant. Lasers Surg. Med. (2019). doi: 10.1002/lsm.23127

27. Davoodzadeh, N. & Cano-Velázquez, M. S. Evaluation of a transparent cranial implant for multi-wavelength intrinsic optical signal imaging. Neural Imaging and (2019).

28. Sakadžić, S. et al. Simultaneous imaging of cerebral partial pressure of oxygen and blood flow during functional activation and cortical spreading depression. Appl. Opt., AO 48, D169–D177 (2009).

29. Bouchard, M. B., Chen, B. R., Burgess, S. A. & Hillman, E. M. C. Ultra-fast multispectral optical imaging of cortical oxygenation, blood flow, and intracellular calcium dynamics. Opt. Express 17, 15670–15678 (2009).

30. Parthasarathy, A. B., Kazmi, S. M. S. & Dunn, A. K. Quantitative imaging of ischemic stroke through thinned skull in mice with Multi Exposure Speckle Imaging. Biomed. Opt. Express 1, 246 (2010).

31. Wang, J., Zhang, Y., Xu, T. H., Luo, Q. M. & Zhu, D. An innovative transparent cranial window based on skull optical clearing. Laser Phys. Lett. 9, 469 (2012).

32. Zhang, C. et al. A large, switchable optical clearing skull window for cerebrovascular imaging. Theranostics 8, 2696–2708 (2018).

33. Zhao, Y.-J. et al. Skull optical clearing window for in vivo imaging of the mouse cortex at synaptic resolution. Light Sci Appl 7, 17153 (2018).

34. Chen, B. R., Bouchard, M. B., McCaslin, A. F. H., Burgess, S. A. & Hillman, E. M. C. High-speed vascular dynamics of the hemodynamic response. Neuroimage 54, 1021–1030 (2011).

35. Heo, C. et al. A soft, transparent, freely accessible cranial window for chronic imaging and electrophysiology. Sci. Rep. 6, 27818 (2016).

36. Smith, S. S., Magnusen, P. & Pletka, B. J. Fracture toughness of glass using the indentation fracture technique. in Fracture Mechanics for Ceramics, Rocks, and Concrete (ASTM International, 1981).

37. Hulbert, S. F. THE USE OF ALUMINA AND ZIRCONIA IN SURGICAL IMPLANTS. in An Introduction to Bioceramics 1, 25–40 (WORLD SCIENTIFIC, 1993).

38. Davoodzadeh, N., Uahengo, G., Halaney, D., Garay, J. E. & Aguilar, G. Influence of low temperature ageing on optical and mechanical properties of transparent yittria stabilized-zirconia cranial prosthesis. in Design and Quality for Biomedical Technologies XI 10486, 104860A (International Society for Optics and Photonics, 2018).

39. Damestani, Y. et al. Transparent nanocrystalline yttria-stabilized-zirconia calvarium prosthesis. Nanomedicine 9, 1135–1138 (2013).

40. Gutierrez, M. I. et al. Novel Cranial Implants of Yttria-Stabilized Zirconia as Acoustic Windows for Ultrasonic Brain Therapy. Adv. Healthc. Mater. 6, (2017).

41. Cano-Velázquez, M. S. et al. Evaluation of Optical Access to the Brain in the Near Infrared Range with a Transparent Cranial Implant. in Latin America Optics and Photonics Conference Tu5C.2 (Optical Society of America, 2018).

42. Davoodzadeh, N., Cuando, N., Aminfar, A. H., Cano, M. & Aguilar, G. ASSESSMENT OF BACTERIA GROWTH UNDER TRANSPARENT NANOCRYSTALLINE YTTRIASTABILIZED-ZIRCONIA CRANIAL IMPLANT USING LASER SPECKLE IMAGING. in LASERS IN SURGERY AND MEDICINE 50, S5–S6 (WILEY 111 RIVER ST, HOBOKEN 07030-5774, NJ USA, 2018).

43. Davoodzadeh, N. et al. Characterization of ageing resistant transparent nanocrystalline yttria-stabilized zirconia implants. J. Biomed. Mater. Res. B Appl. Biomater. (2019). doi: 10.1002/jbm.b.34425

44. Cano-Velázquez, M. S. et al. Enhanced near infrared optical access to the brain with a transparent cranial implant and scalp optical clearing. Biomed. Opt. Express, BOE 10, 3369–3379 (2019).

45. Halaney, D. L. & Jonak, C. OPTICAL COHERENCE TOMOGRAPHY AND LASER SPECKLE IMAGING OF THE BRAIN THROUGH A TRANSPARENT CRANIAL IMPLANT IN A …;. LASERS IN (2018).

46. Davoodzadeh, N. LOW-TEMPERATURE AGEING OF TRANSPARENT NANOCRYSTALLINE YTTRIA-STABILIZED-ZIRCONIA CALVARIUM PROTHESIS. Lasers Surg. Med. (2017).

47. Mishchenko, M. I. V. Tuchin, Tissue Optics: Light Scattering Methods and Instruments for Medical Diagnostics (2nd ed), SPIE Press, Bellingham, WA (2007) Hardbound, ISBN 0-8194-6433-3, xl+841pp. J. Quant. Spectrosc. Radiat. Transf. 110, 528 (2009).

48. Tuchin, V. V. et al. Light propagation in tissues with controlled optical properties. Photon Propagation in Tissues II (1996). doi: 10.1117/12.260832

49. Genina, E. A., Bashkatov, A. N., Sinichkin, Y. P., Yu. Yanina, I. & Tuchin, V. V. Optical Clearing of Tissues: Benefits for Biology, Medical Diagnostics, and Phototherapy. Handbook of Optical Biomedical Diagnostics, Second Edition, Volume 2: Methods doi: 10.1117/3.2219608.ch10

50. Zonios, G., Bykowski, J. & Kollias, N. Skin melanin, hemoglobin, and light scattering properties can be quantitatively assessed in vivo using diffuse reflectance spectroscopy. J. Invest. Dermatol. 117, 1452–1457 (2001).

51. Jacques, S. L. Optical properties of biological tissues: a review. Phys. Med. Biol. 58, R37–61 (2013).

52. Gourley, J. K. & Heistad, D. D. Characteristics of reactive hyperemia in the cerebral circulation. Am. J. Physiol. 246, H52–8 (1984).

53. Zhu, D., Larin, K. V., Luo, Q. & Tuchin, V. V. Recent progress in tissue optical clearing. Laser Photon. Rev. 7, 732–757 (2013).

54. Rutherford, D. et al. Synthesis, characterization, and cytocompatibility of yttria stabilized zirconia nanopowders for creating a window to the brain. J. Biomed. Mater. Res. B Appl. Biomater. (2019). doi: 10.1002/jbm.b.34445

55. Tuchin, V. V. Tissue Optics: Light Scattering Methods and Instruments for Medical Diagnosis. (2015). doi: 10.1117/3.1003040

56. Shi, R. et al. A useful way to develop effective in vivo skin optical clearing agents. J. Biophotonics 10, 887–895 (2017).

57. Alaniz, J. E., Perez-Gutierrez, F. G., Aguilar, G. & Garay, J. E. Optical properties of transparent nanocrystalline yttria stabilized zirconia. Opt. Mater. (2009).

58. Li, P., Ni, S., Zhang, L., Zeng, S. & Luo, Q. Imaging cerebral blood flow through the intact rat skull with temporal laser speckle imaging. Opt. Lett. 31, 1824–1826 (2006).

59. Optical Absorption of Hemoglobin. Available at: https://omlc.org/spectra/hemoglobin/. (Accessed: 13th June 2019)

60. Salehpour, F. et al. Near-infrared photobiomodulation combined with coenzyme Q10 for depression in a mouse model of restraint stress: reduction in oxidative stress, neuroinflammation, and apoptosis. Brain Research Bulletin 144, 213–222 (2019).

61. Caldieraro, M. A. & Cassano, P. Transcranial and systemic photobiomodulation for major depressive disorder: A systematic review of efficacy, tolerability and biological mechanisms. J. Affect. Disord. 243, 262–273 (2019).

62. Hamblin, M. R. Photobiomodulation for traumatic brain injury and stroke. J. Neurosci. Res. 96, 731–743 (2018).

63. Quirk, B. J. et al. Photodynamic therapy (PDT) for malignant brain tumors--where do we stand? Photodiagnosis Photodyn. Ther. 12, 530–544 (2015).

64. Dufour, S. & De Koninck, Y. Optrodes for combined optogenetics and electrophysiology in live animals. Neurophotonics 2, 031205 (2015).

65. Davoodzadeh, N., Cuando, N., Aminfar, A. H., Cano, M. & Aguilar, G. ASSESSMENT OF BACTERIA GROWTH UNDER TRANSPARENT NANOCRYSTALLINE YTTRIASTABILIZED-ZIRCONIA CRANIAL IMPLANT USING LASER SPECKLE IMAGING. in LASERS IN SURGERY AND MEDICINE 50, S5–S6 (WILEY 111 RIVER ST, HOBOKEN 07030-5774, NJ USA, 2018).

66. Richards, L. M. et al. Intraoperative multi-exposure speckle imaging of cerebral blood flow. J. Cereb. Blood Flow Metab. 37, 3097–3109 (2017).

67. Aminfar, A., Davoodzadeh, N., Aguilar, G. & Princevac, M. Application of optical flow algorithms to laser speckle imaging. Microvasc. Res. 122, 52–59 (2019).

68. Zhu, D., Wang, J., Zhi, Z., Wen, X. & Luo, Q. Imaging dermal blood flow through the intact rat skin with an optical clearing method. J. Biomed. Opt. 15, 026008 (2010).

69. Shi, R., Chen, M., Tuchin, V. V. & Zhu, D. Accessing to arteriovenous blood flow dynamics response using combined laser speckle contrast imaging and skin optical clearing. Biomed. Opt. Express 6, 1977–1989 (2015).

70. Ghaffari, H., Grant, S. C., Petzold, L. R. & Harrington, M. G. Regulation of cerebrospinal fluid and brain tissue sodium levels by choroid plexus and brain capillary endothelial cell sodium-potassium pumps during migraine. doi: 10.1101/572727

71. Fowkes, F. G. R. et al. Comparison of global estimates of prevalence and risk factors for peripheral artery disease in 2000 and 2010: a systematic review and analysis. Lancet 382, 1329–1340 (2013).

72. Sokolnicki, L. A., Roberts, S. K., Wilkins, B. W., Basu, A. & Charkoudian, N. Contribution of nitric oxide to cutaneous microvascular dilation in individuals with type 2 diabetes mellitus. Am. J. Physiol. Endocrinol. Metab. 292, E314–8 (2007).

73. Khalil, Z., LoGiudice, D., Khodr, B., Maruff, P. & Masters, C. Impaired Peripheral Endothelial Microvascular Responsiveness in Alzheimer’s Disease. Journal of Alzheimer’s Disease 11, 25–32 (2007).

74. Gnyawali, S. C. et al. High-resolution harmonics ultrasound imaging for non-invasive characterization of wound healing in a pre-clinical swine model. PLoS One 10, e0122327 (2015).

75. Eriksson, S., Nilsson, J. & Sturesson, C. Non-invasive imaging of microcirculation: a technology review. Med. Devices 7, 445–452 (2014).

76. Kisler, K. et al. In vivo imaging and analysis of cerebrovascular hemodynamic responses and tissue oxygenation in the mouse brain. Nat. Protoc. 13, 1377–1402 (2018).

77. Holtmaat, A. et al. Long-term, high-resolution imaging in the mouse neocortex through a chronic cranial window. Nat. Protoc. 4, 1128–1144 (2009).

78. Costantini, I. et al. A versatile clearing agent for multi-modal brain imaging. Sci. Rep. 5, 9808 (2015).

79. Casolco, S. R., Xu, J. & Garay, J. E. Transparent/translucent polycrystalline nanostructured yttria stabilized zirconia with varying colors. Scr. Mater. 58, 516–519 (2008).

80. Filatova, S. A., Shcherbakov, I. A. & Tsvetkov, V. B. Optical properties of animal tissues in the wavelength range from 350 to 2600 nm. J. Biomed. Opt. 22, 35009 (2017).

81. Shi, L., Sordillo, L. A., Rodríguez-Contreras, A. & Alfano, R. Transmission in near-infrared optical windows for deep brain imaging. J. Biophotonics 9, 38–43 (2016).

82. Golovynskyi, S. et al. Optical transparence windows for head tissues in near and short-wave infrared regions. in International Conference on Photonics and Imaging in Biology and Medicine W3A.122 (Optical Society of America, 2017).

83. Cheng, H. et al. Modified laser speckle imaging method with improved spatial resolution. J. Biomed. Opt. 8, 559–564 (2003).

84. Li, N. et al. High spatiotemporal resolution imaging of the neurovascular response to electrical stimulation of rat peripheral trigeminal nerve as revealed by in vivo temporal laser speckle contrast. Journal of Neuroscience Methods 176, 230–236 (2009).

85. Chen, W., Park, K., Volkow, N. D., Pan, Y. & Du, C. Cocaine-Induced Abnormal Cerebral Hemodynamic Responses to Forepaw Stimulation Assessed by Integrated Multi-Wavelength Spectroimaging and Laser Speckle Contrast Imaging. IEEE Journal of Selected Topics in Quantum Electronics 22, 146–153 (2016).

86. Christel, P., Meunier, A., Heller, M., Torre, J. P. & Peille, C. N. Mechanical properties and short-termin vivo evaluation of yttrium-oxide-partially-stabilized zirconia. Journal of Biomedical Materials Research 23, 45–61 (1989).

